# Yolk sac erythromyeloid progenitors sustain erythropoiesis throughout embryonic life

**DOI:** 10.1101/2020.02.27.968230

**Authors:** Francisca Soares-da-Silva, Odile Burlen-Defranoux, Ramy Elsaid, Lorea Iturri, Laina Freyer, Odile Sismeiro, Perpétua Pinto-do-Ó, Elisa Gomez-Perdiguero, Ana Cumano

## Abstract

The first hematopoietic cells are produced in the yolk sac and are thought to be rapidly replaced by the progeny of hematopoietic stem cells. Here we document that hematopoietic stem cells do not contribute significantly to erythrocyte production up until birth. Lineage tracing of yolk sac-derived erythromyeloid progenitors, that also contribute to tissue resident macrophages, shows a progeny of highly proliferative erythroblasts, that after intra embryonic injection, rapidly differentiate. These progenitors, similar to hematopoietic stem cells, are *c-Myb* dependent and are developmentally restricted as they are not found in the bone marrow. We show that erythrocyte progenitors of yolk sac origin require lower concentrations of erythropoietin than their hematopoietic stem cell-derived counterparts for efficient erythrocyte production. Consequently, fetal liver hematopoietic stem cells fail to generate megakaryocyte and erythrocyte progenitors. We propose that large numbers of yolk sac-derived erythrocyte progenitors have a selective advantage and efficiently outcompete hematopoietic stem cell progeny in an environment with limited availability of erythropoietin.

## Introduction

Erythrocytes are the most abundant cells in circulation, they transport oxygen and have a half-life of around 22 days in the mouse, therefore, constant production in the bone marrow (BM) is required to maintain the numbers of circulating red blood cells (RBCs).

Erythropoiesis is the process whereby hematopoietic stem cells (HSC) progressively differentiate into erythro-megakaryocyte and later into lineage-committed erythroid progenitors, immature burst forming unit – erythroid (BFU-E) and the more mature colony-forming unit erythroid (CFU-E). CFU-E successively progress in differentiation through nucleated proerythroblast, basophilic, polychromatophilic and orthochromatic stages, enucleation and formation of RBCs. The distinct stages of erythroid differentiation are characterized by changes in surface expression of the progenitor marker Kit, of the transferrin receptor CD71, of the adhesion molecule CD44 and of the mature erythroid marker Ter119 (Kina *et al.*, 2000; Aisen, 2004; Chen *et al.*, 2009).

During mouse embryogenesis three overlapping hematopoietic waves emerge in distinct anatomic sites. The first blood cells arise in the yolk sac (YS) blood islands at embryonic day (E) 7.5 and belong to the macrophage, erythroid and megakaryocytic lineages (Palis *et al.*, 1999). Primitive erythrocytes are large nucleated cells that express a specific pattern of embryonic (εy- and βH1-) globins (Kingsley *et al.*, 2006). Erythromyeloid progenitors (EMPs) arise in the YS around E8.5 (Bertrand *et al.*, 2005), differentiate into erythrocytes, megakaryocytes, macrophages and other myeloid lineages such as neutrophils, granulocytes and mast cells, but lack HSC activity (Palis *et al.*, 1999; McGrath *et al.*, 2015a, 2015b). EMP-derived erythrocytes resemble definitive erythrocytes and express embryonic βH1- and adult β1- but no εy-globins (McGrath *et al.*, 2011). Immature HSC (imHSC) emerge after E8.5 (E8.5-E11.5) (Cumano *et al.*, 1996; de Bruijn *et al.*, 2000; Taoudi *et al.*, 2008; Bertrand *et al.*, 2010; Kissa and Herbomel, 2010; Kieusseian *et al.*, 2012) in the major arteries, through an endothelial to hematopoietic transition process, rapidly enter circulation and colonize the fetal liver (FL) where they expand and differentiate, generating the blood lineages. EMP that arise through a similar process in the YS (Frame *et al.*, 2016; Kasaai *et al.*, 2017) also converge to the FL where they are identified as Kit^+^CD16/32^+^ in contrast to Kit^+^CD16/32^-^ imHSC.

The analysis of *c-Myb* mutants, where primitive hematopoiesis is preserved but HSC-derived hematopoiesis is lacking, indicated that YS-derived primitive hematopoietic cells sustain embryonic life up until E15.5 (Mucenski *et al.*, 1991; Tober *et al.*, 2008; Schulz *et al.*, 2012). Mice mutant for the Runx1 partner CBFβ have impaired EMP and HSC formation and lack long-term reconstitution activity (Wang *et al.*, 1996). Selective expression of CBFβ in Tek or Ly6a expressing cells results in the rescue of YS EMP or HSC. In the absence of HSC activity, EMP-derived hematopoietic cells maintain viable embryos throughout development up until birth (Chen *et al.*, 2011).

YS hematopoiesis has long been considered a transient wave devoted to the production of erythrocytes, megakaryocytes and a few myeloid cells that ensure oxygenation and tissue hemostasis. Thus, HSCs-derived hematopoiesis was thought to replace YS-derived cells shortly after HSC migrate to the FL at E10.5 (Palis, 2016). Recently, however, growing evidence endows the YS with the capacity to contribute to tissue resident cells such as macrophages that persist throughout life (Gomez Perdiguero *et al.*, 2015) and mast cells (Gentek *et al.*, 2018) maintained up until birth. Primitive erythrocytes were also shown to persist throughout gestation (Fraser *et al.*, 2007) and EMP-derived cells contribute to the erythrocyte compartment for more than 20 days upon transplantation (McGrath *et al.*, 2015a). Nonetheless, it has been difficult to establish the temporal relative contribution of EMP or HSC-derived progenitors to erythropoiesis because they share surface markers and transcriptional regulators and are therefore indistinguishable.

Here we report a large population of Kit^+^CD45^-^Ter119^-^ erythroid progenitors unique to FL, comprising >70% of E14.5 Ter119^-^CD45^-^ cells (>10% of FL cells). These are the most actively proliferating progenitors at early stages and progress in erythroid differentiation through the upregulation of the surface marker CD24 concomitant with that of CD71, with subsequent loss of Kit and upregulation of Ter119. These cells that require *c-Myb* expression originate from YS EMP as they are co-labeled with microglia in the Csf1r^MeriCreMer^Rosa26^YFP^ lineage-tracer model. They persist through fetal life and are the major contributors to the RBC compartment. In a lineage tracer model, we show that Flt3 expressing progenitors that comprise most HSC progeny do not contribute significantly to embryonic erythropoiesis. HSC erythroid progenitors require higher concentrations of erythropoietin (Epo) than their YS-derived counterparts, for erythrocyte differentiation. The limiting amounts of Epo available in the embryo results in a selective advantage of YS-derived over HSC-derived erythropoiesis.

## Results

### A unique population of Kit^+^ cells represents the majority of FL Ter119^-^CD45^-^ cells

We analyzed, by flow cytometry, Kit expression in the FL together with antibodies that identify erythrocytes, hematopoietic, endothelial and epithelial cells at different embryonic time-points. We detected a large fraction of Kit^+^ cells (>50%) expressing neither Ter119 nor CD45 (Fig. 1A, Supplementary Fig. 1A). Single-cell surface marker expression data from E14.5 FL cells was projected as tSNE1 vs tSNE2 (Fig. 1A) and three major clusters were defined by the expression of epithelial cadherin (E-Cadherin/CD324) on epithelial cells, platelet/endothelial cell adhesion protein (PECAM-1/CD31) on endothelial cells and Kit. Combined analysis of Kit expression together with CD24 frequently associated with immature hematopoietic cells further defined 3 populations in the Ter119^-^CD45^-^CD31^-^CD324^-^ compartment: Kit^+^CD24^-^ (hereafter called P1), Kit^+^CD24^+^ (P2) and CD24^+^Kit^-^ (P3) cells (Fig. 1B; Supplementary Fig. 1A). P1 cells decreased while P2, and subsequently P3 cells increased as gestation progressed (Fig. 1C). Numbers of P2 cells reached a maximum (around 10^6^ cells per FL) at E14-15 and decreased thereafter, although they were still detected around birth (E18.5) (Fig. 1D). Kit^+^CD45^-^Ter119^-^Lin^-^ (P1 and P2) cells were are also negative for the expression of Sca-1 that marks multipotent progenitors, for CD16/32 that marks GM progenitors and for CD34 marking common myeloid progenitors (CMP) and therefore they fall in a gate that typically defines megakaryocyte/erythrocyte progenitors (MEP) in the FL and in the BM (Supplementary Fig. 1B). Unlike their BM counterparts however, where all Kit^+^ cells co-expressed CD45, most FL Kit^+^ (P1 and P2) cells within the LK compartment did not express CD45 (Fig. 1E and 1F) raising the possibility that they did not belong to the hematopoietic lineage. We therefore identified a major population of Kit^+^CD45^-^Ter119^-^ cells unique to FL of undefined lineage affiliation and origin.

**Figure 1.**
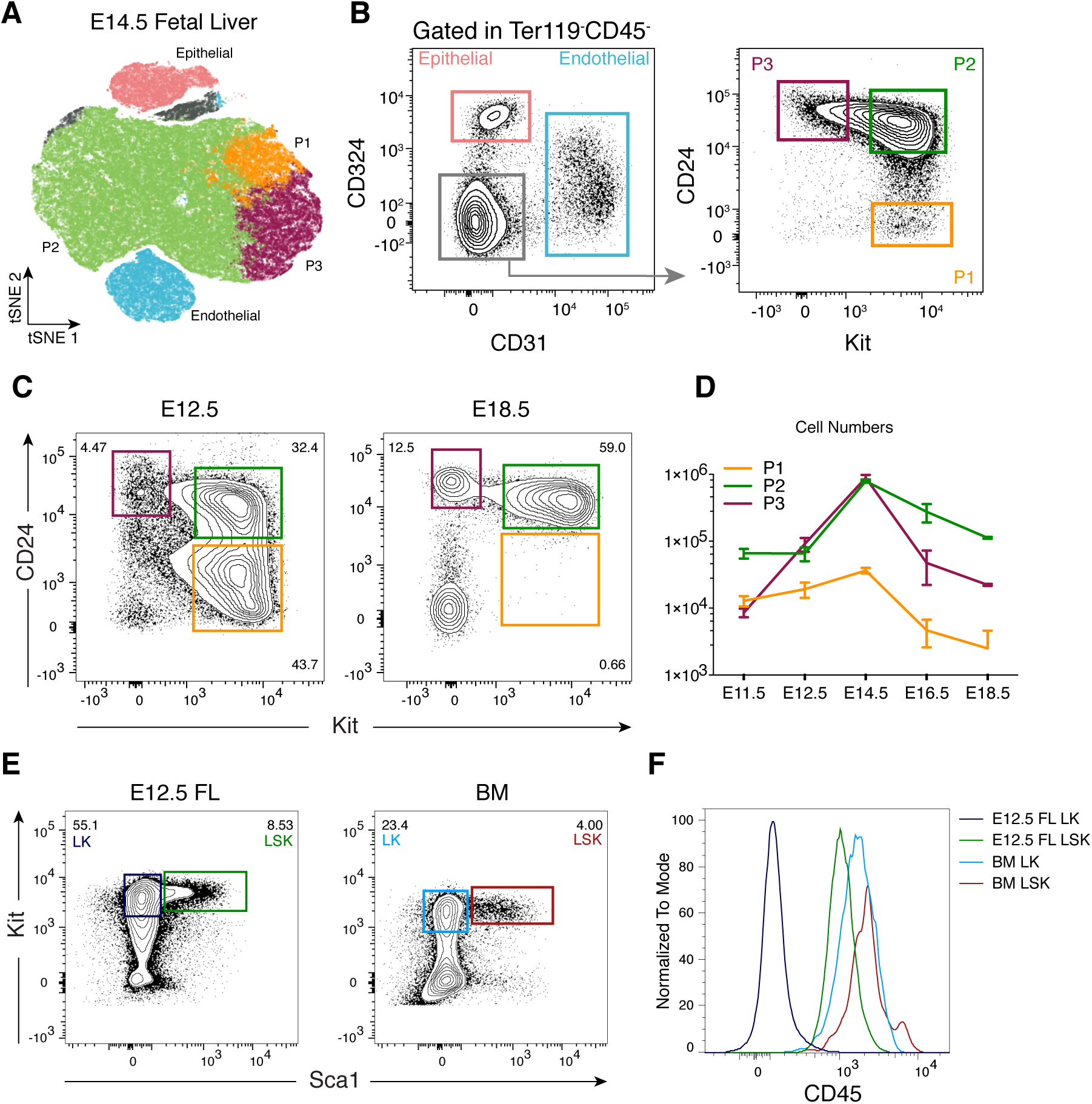
A population of Kit^+^ cells unique to FL represents the majority of Ter119^-^CD45^-^ cells. **(A)** tSNE analysis and hierarchical clustering of flow cytometry data of Ter119^-^CD45^-^ cells from E14.5 fetal livers stained with the surface markers CD31 (endothelial cells), CD324 (epithelial cells, hepatoblasts), Kit and CD24. **(B)** Representative FACS plots of the clusters identified in **(A)** using the same color code. **(C)** Representative FACS plots of TER119^-^CD45^-^ cells from E12.5 and E18.5 regarding Kit and CD24 expression and **(D)** corresponding cell numbers at different stages. **(E)** Phenotype of viable Lin^-^ (the lineage cocktail contains Ter119, Gr-1, CD19, CD3, CD4, CD8, Nk1.1, Il7r, TCRαβ, TCRγδ and Cd11c for BM and Ter119, Gr-1, CD19, Nk1.1, Il7r for E12.5 fetal liver) E12.5 FL and BM cells using Kit and Sca1. **(F)** Histogram of CD45 expression in Lin^-^Kit^+^Sca1^-^ (LK) and Lin^-^Kit^+^Sca1^+^ (LSK) cells from E12.5 FL and adult BM. (E11.5 n= 6, E12.5 n=17, E14.5 n=4, E16.5 n=8, E18.5 n=2). See also Supplementary Fig. 1.

### P1 and P2 cells in the FL have an erythroid progenitor signature

To investigate the cellular identity of Kit^+^CD45^-^Ter119^-^ FL cells we performed RNA sequencing of the three major populations P2, CD324^+^ and CD31^+^ cells. Within the highest expressed transcripts in P2 cells were *Myb*, *Bcl11a*, *Klf1*, *Gata1* and *Epor*, hematopoietic specific transcripts most of which are associated with erythrocyte differentiation (Fig. 2A). The 122 genes upregulated >2-fold in P2 vs CD324^+^ cells were subjected to gene ontology analysis using *Enrichr* (Chen *et al.*, 2013) (List of submitted genes in Supplementary Table 1). The top biological processes and tissue associated genes revealed an erythrocyte/erythroblast profile (Fig. 2B). These results were validated by Q-RT-PCR indicating that *Gata1*, *Lmo2*, *Klf1* and *Epor* expressions gradually increased from P1 to P3 cells, with the latter showing comparable expression levels of these transcripts to Ter119^+^ erythroblasts (Fig. 2C). Hemoglobin transcripts for *Hbb-y*, *Hbb-bh1* and *Hbb-b1* were detected in P3 cells but only significantly expressed in Ter119^+^ cells. Multipotent hematopoietic associated transcription factors such as *c-Myb*, *Runx3* and *Bmi1* decreased as the erythroid specific transcripts increased. The results above indicated that the CD45^-^ subsets (P1 and P2) identified by the expression of Kit and CD24 (also found on all erythrocytes), are erythroid progenitors and suggested a hierarchy where immature P1 cells further differentiate into P2 and later lose Kit expression (P3) before acquiring Ter119 expression, the definitive marker of erythroid identity.

**Figure 2.**
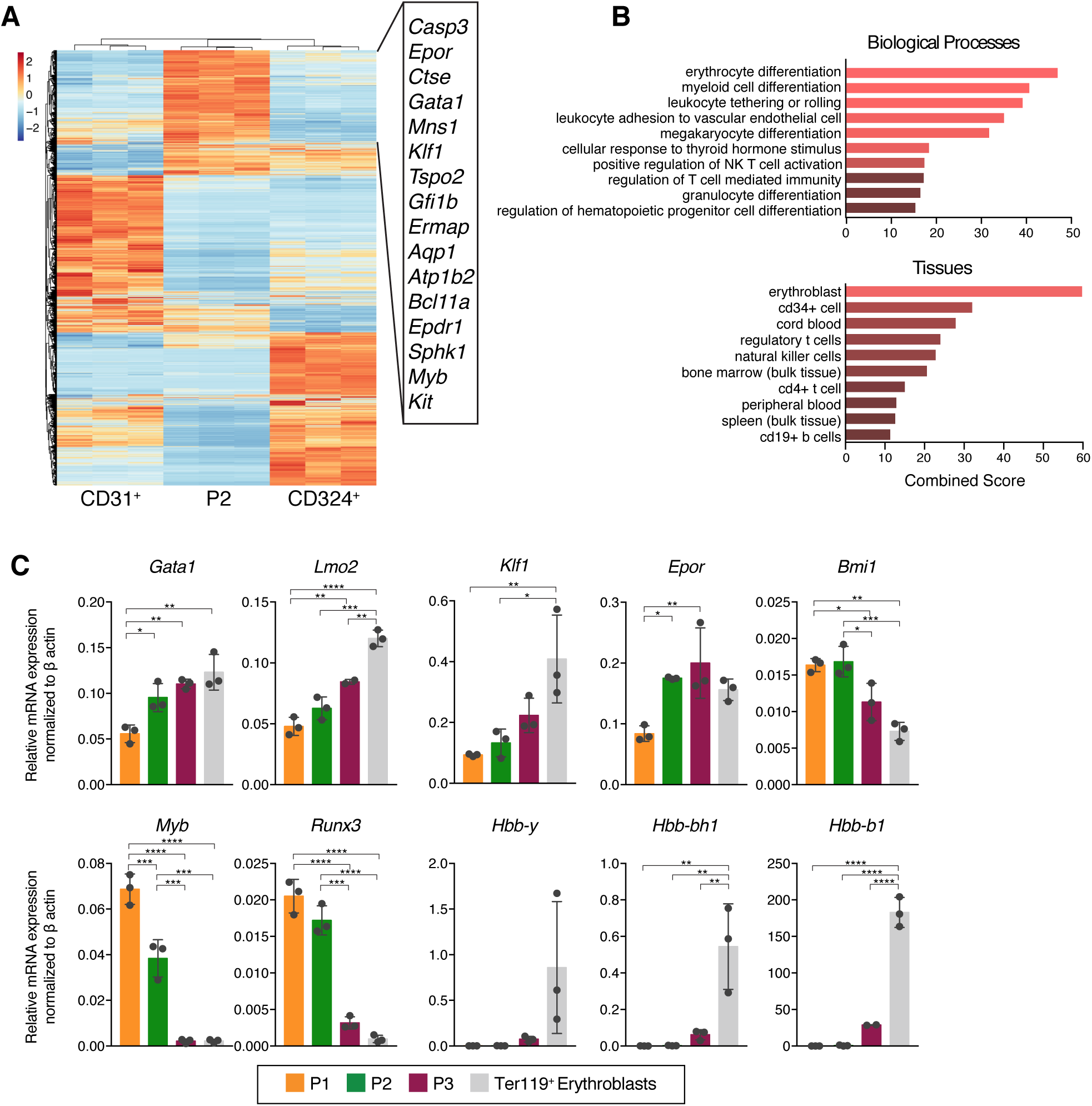
P1 and P2 cells in the FL have an erythroid progenitor transcriptional signature. **(A)** CD31^+^, P2 and CD324 ^+^ cells from E14.5 FL were sorted and subjected to RNA sequencing. Differentially expressed genes are represented as heatmap and most expressed genes are listed (n=3 independent litters). **(B)** Gene set enrichment analysis on genes with more than two-fold difference in expression level between P2 and CD324 ^+^ cells using *Enrichr* web-application; top 10 significantly associated GO Biological Processes and ARCHS4 Tissues are shown. Top biological process is erythrocyte differentiation (q-value <1.0e^-6^; Gene Ontology term GO:0030218) and top tissue associated is erythroblast (q-value <1.0e^-14^; ARCHS4 Tissues gene database). Gene lists available on request. **(C)** E14.5 FL P1, P2, P3 and Ter119^+^ cells were sorted and gene expression of key erythroid genes (Gata1, Lmo2, Klf1, Epor and Bmi1), progenitor associated genes (*c-Myb* and *Runx3*) and hemoglobins (Hbb-y, Hbb-bh1, Hbb-b1) was analyzed. QPCR data was analyzed using the delta Ct method and was normalized with β-actin. Statistical significance was assessed using one-way ANOVA followed by Tukey’s multiple comparison test **(C)** *p<0.05, **p<0.01, ***p<0.001, ****p<0.0001. Data are represented as mean ± standard deviation from 3 independent experiments.

### P1, P2 and P3 cells represent increasingly mature stages within the erythroid lineage

Erythroid differentiation has been characterized by the expression of CD71 (transferrin receptor) and CD44, in Ter119^+^ cells (McGrath *et al.*, 2017). Imaging flow cytometry (Fig. 3A) showed that the proerythroblast marker CD71 was low in P1 but expressed in P2 and highly expressed in all P3 cells (Fig. 3B), indicating they correspond to consecutive stages of erythrocyte development.

**Figure 3.**
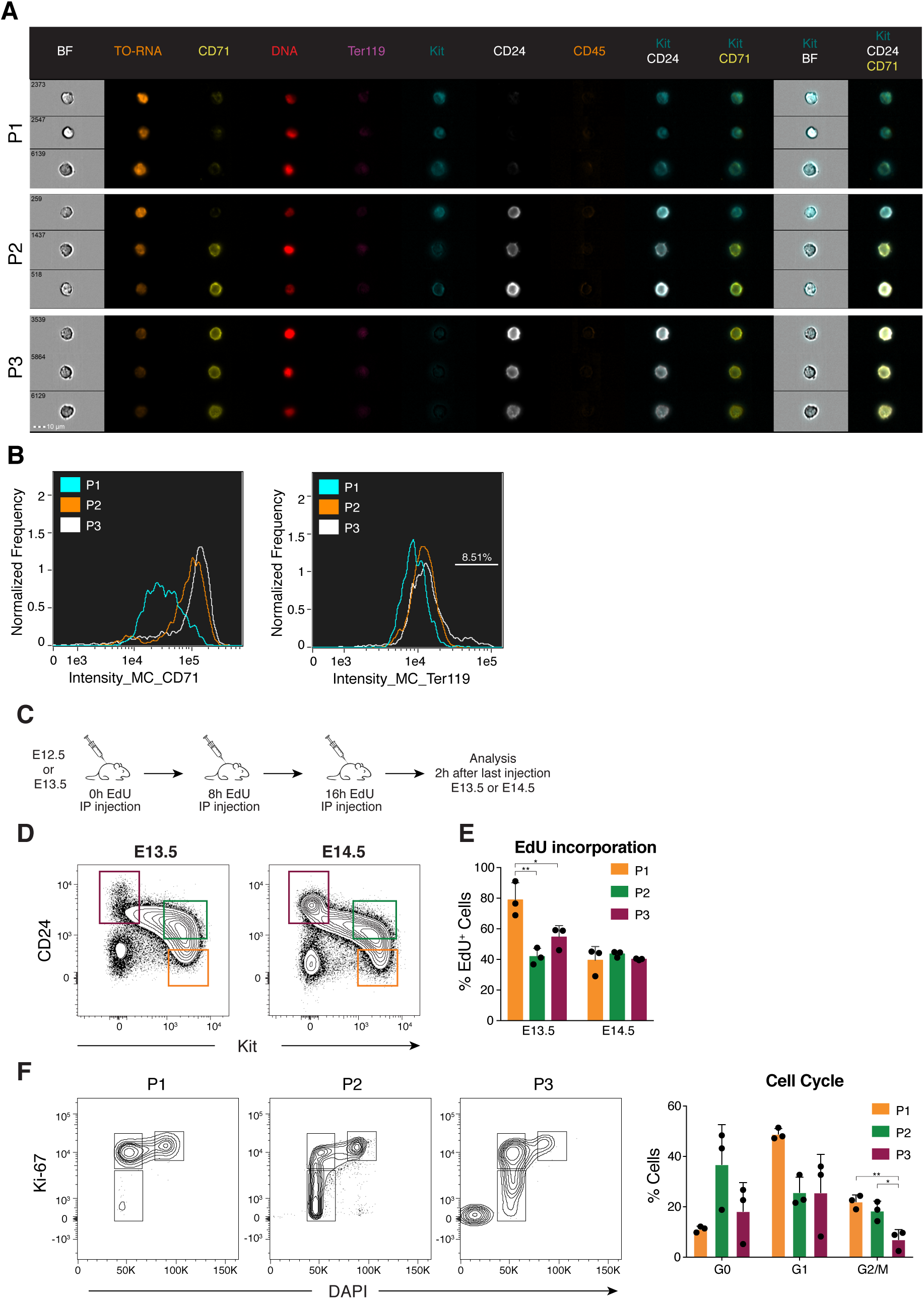
P1, P2 and P3 cells represent increasingly maturation stages within the erythroid lineage. **(A)** E13.5 Ter119^-^ CD45^-^ cells were analyzed by imaging flow cytometry using CD71, Ter119, Kit, CD24 and CD45 as surface markers, DRAQ5 to label nuclei and Thiazole Orange (TO) to label RNA. Representative images of P1, P2 and P3 cells. **(B)** Expression of CD71 and Ter119 was assessed in P1, P2 and P3 cells and plotted as a histogram. **(C)** Experimental design of cell cycle analysis using EdU labelling. E12.5 or E13.5 pregnant mice were injected intraperitoneally with 100 ug of EdU at 0h, 8h and 16h. Fetal livers were collected 2h after the last injection and EdU expression was analyzed on Ter119^-^CD45^-^CD54^-^CD31^-^ cells using the indicated gates **(D)**. **(E)** Percentages of EdU incorporation in P1, P2 and P3 cells at E13.5 and E14.5 (n=3). **(F)** Cell cycle analysis of E14.5 fetal liver cells using Ki-67 and DAPI. Statistical significance was assessed using one-way ANOVA followed by Tukey’s multiple comparison test **(C)** *p<0.05, **p<0.01, ***p<0.001, ****p<0.0001. Data are represented as mean ± standard deviation from 3 independent experiments.

In BM and FL, progenitors that form small erythroid colonies (CFU-E) are characterized as Kit^+^CD71^+^Ter119^-^ whereas low levels of Ter119 expression marks proerythroblasts that lost proliferative capacity (McGrath *et al.*, 2017). P2 FL cells express CD71 in the virtual absence of Ter119 indicating that they correspond to CFU-E. P3 cells express low levels of Ter119 (Fig. 3B) visible in 8% of them and express erythroid genes at levels similar to Ter119^+^ erythroblasts (Fig. 2C) suggesting they correspond to proerythroblasts.

CD71 expression is limited to P2 and P3 cells indicating that CD71 and CD24 are redundant markers in this context, further confirmed by conventional flow cytometry (Supplementary Fig. 1C).

To examine the cell cycle status of the 3 cell subsets we injected timed pregnant females with the nucleotide analog 5-ethynyl-2′-deoxyuridine (EdU) to label newly synthesized DNA. E12.5 or E13.5 pregnant females were given 3 EdU injections with 8h intervals and FLs were analyzed 2h after the last injection (Fig. 3C). 80% of P1 E13.5 FL cells were labelled with EdU compared to around 50% of P2 and P3 cells. E14.5 FL cells show the same percentage of EdU incorporation (40-50%) in all 3 subsets (Fig. 3D and 3E). Cell proliferation was further assessed by analyzing the expression of the nuclear protein Ki-67 that in association with DAPI allows distinguishing cells in G0, G1 and G2M phases of the cell cycle. Consistent with the EdU labelling experiments, P1 showed the lowest frequency of cells in G0 (∼10%) and the highest frequency of cells expressing Ki-67, from which around 20% are actively synthesizing DNA (DAPI^+^) (Fig. 3F). By contrast, P3 cells show the lowest frequency of proliferating cells (Fig. 3F). Taken together these results indicated that P1 are the most proliferating cells and, as they transit onto the P2 and further into the P3 subset, lose proliferative activity.

### P2 and P3 cells are committed erythroid progenitors whereas P1 retain residual myeloid potential

To assess the differentiation potential of CD45^-^Kit^+^ FL cells we performed differentiation assays in liquid cultures and in semi-solid colony assays (Fig. 4A and 4B). E13.5 P1, P2, P3 and Lin^-^CD45^+^Kit^+^Sca1^-^ (LK) cells, as control, were sorted and cultured in the presence of Scf, Epo, Tpo, M-CSF and GM-CSF to allow differentiation into erythroid, megakaryocyte and myeloid lineages. Limiting-dilution analysis showed that P1 and P2 gave rise to hematopoietic colonies at frequencies similar to that of LK cells (1 in 1 for LK and P1, 1 in 2 for P2 cells) while P3 cells did not divide significantly in culture (less than 1 colony in 2592 wells analyzed) (Fig. 4A). In colony assays both P1 and P2 cells gave a majority of CFU-E (more than 50% of plated cells). P1 cells generated also BFU-E (<5%), CFU-M (5%) and CFU-Mk (5%) whereas control (LK) generated a majority of myeloid colonies (CFU-G, CFU-GM and CFU-GEMM) and less than 5% of CFU-E (Fig. 4B).

**Figure 4.**
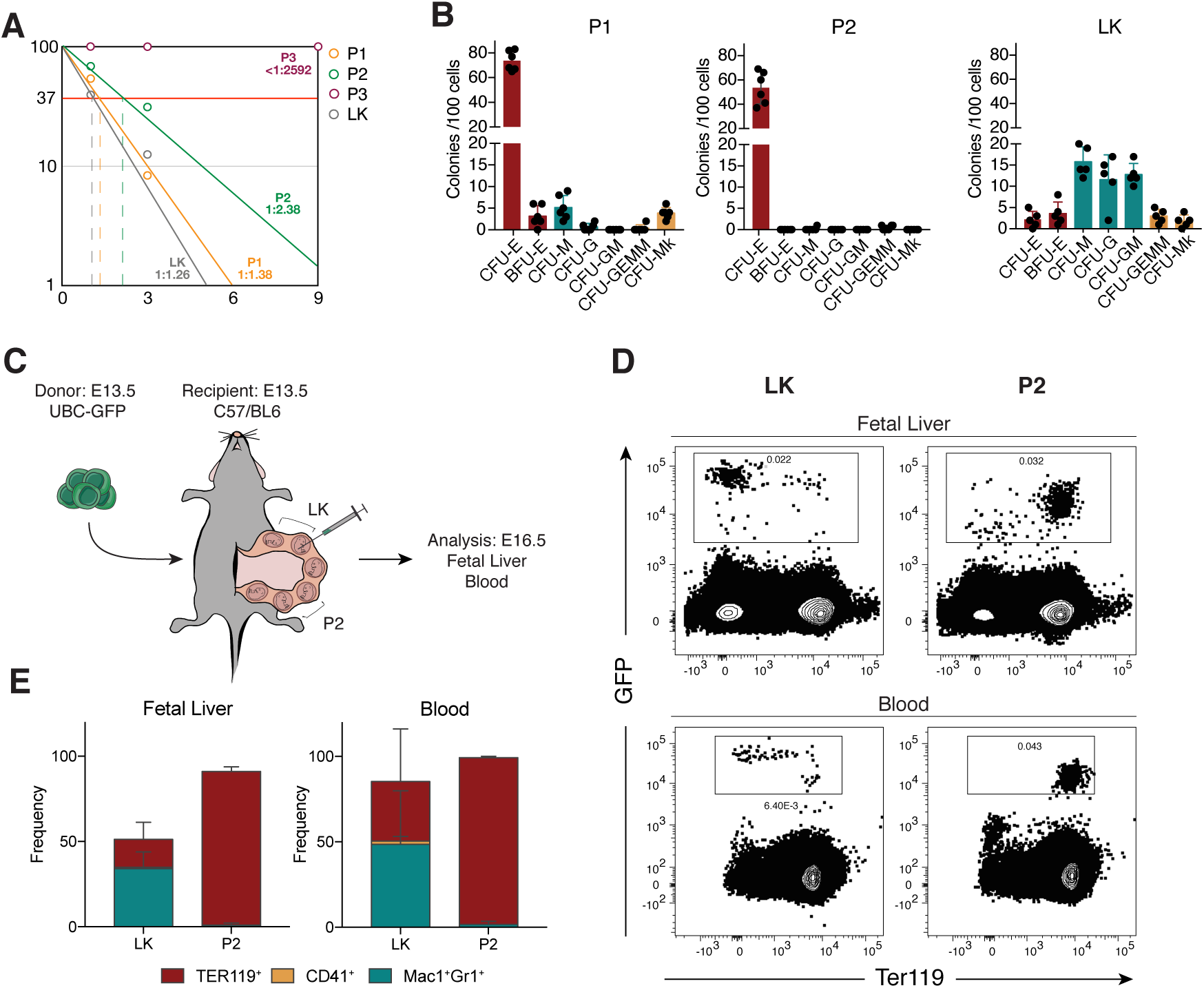
P2 and P3 cells are committed erythroid progenitors whereas P1 retain residual myeloid potential. **(A)** Frequencies of colony forming cells in P1, P2, P3 and Lin^-^ CD45^+^ Kit^+^ (LK) cells from E13.5 FL (n= 288 wells from 3 independent experiments of each population). **(B)** *In vitro* lineage potential of E13.5 P1, P2, P3 and LK cells in semi-solid cultures. CFU-E colonies were quantified at 3 days and BFU-E, CFU-M, CFU-G, CFU-GM, CFU-GEMM and CFU-Mk at 7 days of culture (n= 6, 3 independent experiments). **(C)** Schematic representation of transplantation experiment. E13.5 C57/BL6 pregnant females were anesthetized, the peritoneal cavity was opened, and the uterus exposed. Embryos were injected intraperitoneally with 20000 E13.5 GFP^+^ purified cells from UBC-GFP embryos. FL and blood were collected 3 days post-injection. **(D)** Representative FACS plots showing erythroid contribution of GFP^+^ cells in fetal liver and blood after injection of P2 or LK cells and quantification **(E)** (P2 n=3, LK n=4, 4 independent experiments). Data are represented as mean ± standard deviation.

We then probed the differentiation potential of P2 cells *in vivo*. Cells purified from E13.5 UBC-GFP embryos were injected into E13.5 C57/BL6 recipient embryos, *in utero* (Fig. 4C). FL and blood collected three days later indicated that GFP^+^P2 originated exclusively GFP^+^Ter119^+^ cells whereas LK generated a majority of myeloid cells detected both in FL and in blood (Fig. 4D and 4E) while none gave rise to lymphocytes.

These results demonstrated that P2 FL cells are committed erythroid progenitors while P1 cells retain residual *in vitro* myeloid differentiation potential.

### P1/P2 progenitors require *c-Myb* expression

The transcription factor *c-Myb* is essential for adult hematopoieisis. Myb^-/-^ embryos have impaired definitive hematopoiesis that affects all lineages and are not viable after E15. Only primitive YS-derived erythropoiesis, primitive megakaryocytes (Tober *et al.*, 2008) and tissue-resident macrophages (Schulz *et al.*, 2012) have been described in *c-Myb* mutants. To test whether P1/P2 FL cells were affected by the *c-Myb* mutation we analyzed Myb^+/-^ and Myb^-/-^ E14.5 embryos. Ter119^+^ cells were drastically decreased in frequency and numbers in Myb^-/-^ FLs when compared to heterozygous littermates (Fig. 5A and 5B). P1 cells were undetectable in Myb^-/-^ while P3 cells were present, albeit in reduced numbers (Fig. 5B). *c-Myb* was reported to regulate c-Kit expression (Ratajczak *et al.*, 1998) and because we failed to detect Kit^+^ (P1 and P2) cells in the Myb^-/-^ FL we considered the possibility that erythroid progenitors, although unable to express Kit, might be present in mutant FL. We sorted E14.5 CD24^-^, CD24^+^ and Ter119^+^ cells from Myb^+/-^ and Myb^-/-^ FLs and analyzed the expression of erythroid genes. *Epor*, *Tal1* and *Klf1* were not detected in CD24^-^ and CD24^+^Myb^-/-^, compared to Myb^+/-^ cells (Fig. 5C). Only Ter119^+^ cells expressed detectable levels of *Epor* and *Tal1* together with high levels of *Hbb-y* indicating they represent primitive erythrocytes. Of note, *Klf1*, a transcription activator of the β-globin promoter, was not expressed in primitive Ter119^+^Myb^-/-^ cells (Fig. 5C). These results demonstrated that differentiation and/or survival of CD45^-^Kit^+^ erythroid progenitors required the transcription factor *c-Myb*.

**Figure 5.**
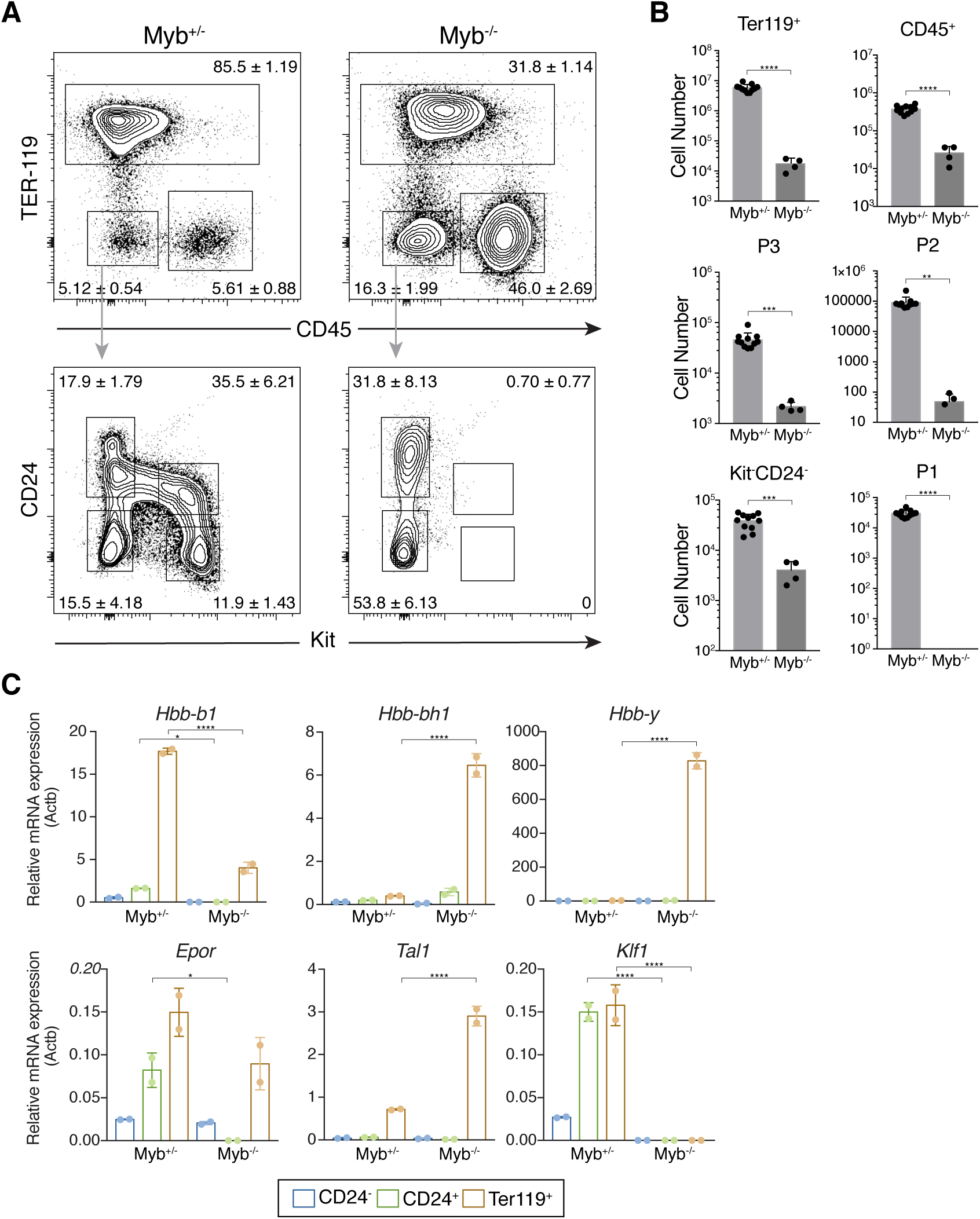
P1/P2 progenitors require c-Myb expression. **(A)** Representative FACS plots showing percentages of Ter119 and CD45 (top panel) and Kit and CD24 (lower panel) expressing cells in *c-Myb*^+/-^ and *c-Myb*^-/-^ E14.5 FL and corresponding absolute numbers **(B)**. **(C)** Expression of hemoglobin (*Hbb-b1*, *Hbb-bh1* and *Hbb-y*) and key erythroid genes (*Epor*, *Tal1* and *Klf1*) in CD24^-^, CD24^+^ and Ter119^+^ cells were analyzed by qPCR. Statistical significance was assessed using one-way ANOVA followed by Tukey’s multiple comparison test *p<0.05, **p<0.01, ***p<0.001, ****p<0.0001. Data are represented as mean ± standard deviation from 2 independent experiments.

### P1/P2 cells originated from YS progenitors and were major contributors to embryonic erythropoiesis

P1/P2 (CD45^-^Kit^+^) cells were not detected in the adult BM, suggesting that they represent a transient hematopoietic population. To assess the origin of P1/P2 FL cells we analyzed FL cells from Csf1r^MeriCreMer^Rosa26^YFP^ pregnant females pulsed with a single dose of OH-TAM at E8.5 (Fig. 6A). Induction of the Cre recombinase at this developmental stage marks tissue resident macrophages but virtually no HSC-derived progenitors (Gomez Perdiguero *et al.*, 2015). At E11.5, P1 and P2 cells were labelled to comparable levels of those in the microglia, taken as positive control (Fig. 6B). In subsequent days the frequency of YFP labelled P3 and erythroblast (Lin^+^CD71^+^) cells, that represent more differentiated erythroid subsets than P2, increased whereas that of more immature P1 decreased. In line with previous reports YFP labelled LSK cells were undetectable (Fig. 6B, Supplementary Fig. 2). The dynamic of YFP labelled erythroid progenitors is consistent with a progression in erythroid differentiation and indicates a lineage relationship between the 3 subsets. The frequency of YFP labelled P1 and P2 decreases between E11.5 and E13.5, a dynamic that is best explained by a fast differentiation progression.

**Figure 6.**
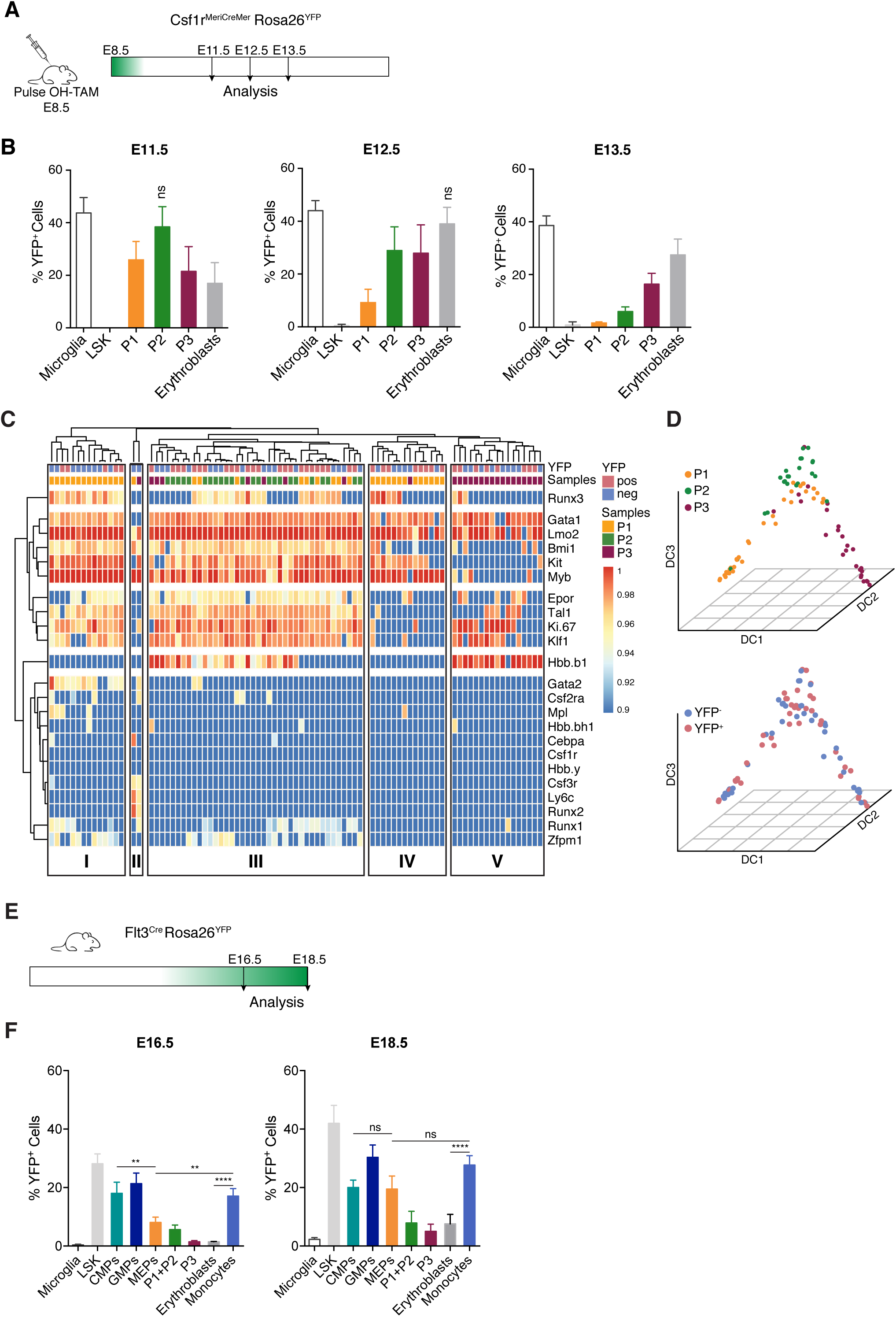
P1/P2 cells originated from yolk sac and were major contributors to embryonic erythropoiesis. **(A)** Experimental design for lineage tracing experiments using the Csf1r^MeriCreMer^Rosa26^YFP^. Cre recombinase expression was induced in Csf1r^MeriCreMer^Rosa26^YFP^ pregnant females with a single injection of OH-TAM at E8.5 and embryos were analyzed 3, 4 and 5 days after injection. **(B)** Percentage of YFP+ cells at E11.5, E12.5 and E13.5 after a pulse of OH-TAM at E8.5 (E11.5 n=12, E12.5 n=11, E13.5 n=7). **(C)** Heatmap of single-cell qPCR in sorted cells from Csf1r^MeriCreMer^ E13.5 FL pulsed with OH-TAM at E8.5. Each column represents a single-cell and its color-coded according to YFP expression (YFP^+^ vs YFP^-^) and cell type (P1, P2, P3). Gene expression was normalized to *b-actin* and *Gapdh* and unsupervised hierarchical clustering was performed. **(D)** Diffusion map of the 3 populations based on single-cell gene expression. Left panel is color-coded according to cell type and right panel according to YFP expression. (A total of 87 cells were analyzed). **(E)** Experimental design for lineage tracing experiments using the Flt3^Cre^Rosa26^YFP^ mice model. Embryos were analyzed at E16.5 and E18.5. **(F)** Frequency of YFP expressing cells at E16.5 and E18.5 (E16.5 n=3, E18.5 n=4) in CMPs (Lin^-^c-kit^+^Sca-1^-^Flt3^-^CD71^-^ CD16/32^-^CD34^+^), GMPs (Lin^-^c-kit^+^Sca-1^-^Flt3^-^CD71^-^CD16/32^+^CD34^+^), MEPs (Lin^-^c-kit^+^Sca-1^-^Flt3^-^CD71^-^CD16/32^-^CD34^-^), P1+P2 (Lin^-^CD45^-^c-kit^+^), P3 (Lin^-^CD45^-^Kit^-^ CD71^+^), erythroblasts (Lin^+^CD71^+^) and monocytes (Lin^-^CD45^+^CD11b^int^F4/80^-^) cells. Microglia (CD45^+^F4/80^+^CD11b^+^) and LSK (Lin^-^CD45^+^Sca1^+^Kit^+^) cells were used as controls. See also Supplementary Figs. 2 and 3.

We considered the possibility that YFP^+^ and YFP^-^ CD45^-^Kit^+^ FL cells represented two distinct populations. To test for this hypothesis, we sorted YFP positive and negative P1, P2 and P3 cells from E13.5 FL pulsed at E8.5 and performed single cell multiplex gene expression analysis for progenitor, erythroid and myeloid genes. Unsupervised hierarchical clustering did not segregate YFP^+^ from YFP^-^ cells indicating that they have a similar transcriptional profile and therefore likely do not represent two divergent progenitor populations (Fig. 6C). Cluster I and IV contained P1 cells characterized by the expression of *Gata1*, *Lmo2* and *c-Myb*. Cluster I differed from Cluster IV by high frequency of cells expressing *Epor*, *Tal1* and *Klf1*. Interestingly, few cells in this cluster also co-expressed the myeloid factors *Runx1*, *Gata2*, *Zfpm1* and *Mpl*, suggesting a broad myeloid transcriptional priming, consistent with data from *in vitro* differentiation assays (Fig. 4B). Few cells segregated from all other in Cluster II defined by expression of *Csf3r*, *Ly6c* and *Runx2* in the absence of erythroid associated transcripts. Cluster III comprises a majority of P2 cells, expressing high levels of erythroid genes and low levels of hemoglobin, thus defining a transitional erythroid population. Cluster V contains P3 cells that express high levels of hemoglobin in the absence of *c-Kit* or *c-Myb*.

To analyze the differentiation trajectory between the 3 populations we generated a diffusion map and according with the previous observations obtained a trajectory in which P1 cells progress through a P2 stage and subsequently a P3 stage (Fig. 6D). This differentiation trajectory is in line with the gene expression data (Fig. 2C), with the imaging flow cytometry results (Fig. 3A) and with the clonal differentiation assays (Fig. 4B). YFP^+^ and YFP^-^ cells do not show distinct trajectories indicating they do not represent different progenitor subsets (Fig. 6D).

### HSC do not contribute to erythropoiesis up until birth

To evaluate HSC contribution to fetal erythropoiesis we analyzed Flt3^Cre^Rosa26^YFP^ embryos where multipotent progenitors (MPP Flt3^+^) of HSC origin and their progeny express YFP (Benz *et al.*, 2008; Buza-Vidas *et al.*, 2011) in addition to a transient population of Flt3^+^ progenitors recently identified (Beaudin *et al.*, 2016) (Fig. 6E). Less than 2% of microglial cells expressed YFP at any given time-point analyzed. LSK were increasingly labelled with >30% of YFP^+^ cells at E16.5, reaching >40% at E18.5 (Fig. 6F). Both CMP and GMP compartments exhibited similar levels of YFP expression to those in LSK, and the same was observed for Cd11b^+^F4/80^-^ monocytes, consistent with their definitive HSC origin. By contrast, only less than 5% P1+P2 or P3 were labelled with YFP by E16.5 and less than 10% by E18.5. Erythroblasts showed less than 8% of YFP labelled cells at all analyzed time-points indicating that HSC are not contributing to mature erythrocytes up until E18.5, the day before birth. MEP showed a delayed labelling profile with 8% at E16.5 reaching 20% around birth. Taken together these observations indicated that neither HSC-derived nor any other Flt3 expressing progenitor contribute significantly to erythropoiesis throughout fetal life and that the erythrocyte progenitors that sustain fetal erythropoiesis differentiate from YS-derived EMPs.

FL stroma produces Epo, essential for erythrocyte production, albeit at lower concentrations than kidney, the adult source of Epo (Suzuki *et al.*, 2011). Embryonic progenitors have been reported to react to lower concentrations of Epo than their adult counterparts (Rich and Kubanek, 1980). We compared erythrocyte production from P2 of YS origin with that from CD45^+^ MEP of HSC origin from the same FL (Supplementary Fig. 4). HSC-derived MEP were about two-fold less efficient in generating erythrocytes than YS-derived P2 at limiting levels of Epo and required more than ten-fold higher concentrations to reach 50% of erythrocyte colony formation. These results provide experimental evidence for the mechanism controlling the selection of YS-derived over HSC-derived progenitors in fetal erythropoiesis.

## Discussion

Here we describe a population of CD45^-^Kit^+^ (P1/P2) hematopoietic cells unique to FL, found from E11.5 up until birth and that, at its peak (around E14.5), represent a major population comprising more than 70% of the CD45^-^Ter119^-^ FL cells and 10-15 % of total FL cells. *In vitro* analysis of their differentiation potential showed that they give rise to erythroid colonies (50-70%) at higher frequencies than adult BM MEPs (∼15%) (Akashi *et al.*, 2000). The majority of CD45^-^Kit^+^ cells express CD24, a marker found in immature hematopoietic and other cell types and mature erythrocytes, and up-regulated concomitantly with CD71, the transferrin receptor, a marker only found in erythroid progenitors (Dong *et al.*, 2011).

Gene expression analysis and *in vitro* assays indicated a developmental progression where P1 cells further develop into P2 and later into P3 cells before acquisition of Ter119 expression. Both P1 and P2 cells contained high frequency of CFU-E and had increasing expression of the erythroid genes *Gata1* and *Epor*. By contrast, P3 cells did not generate colonies *in vitro* and expressed the erythroid transcripts *Gata1*, *Lmo2*, *Klf1* and *Epor* at levels similar to Ter119^+^ cells, stage at which they also express *Hbb-b1*, low levels *of Hbb-bh1* and undetectable *Hbb-y* transcripts, indicating they are within the definitive erythroid lineage. Of notice, P1 cells are among the most actively proliferative FL progenitors indicating that they can considerably expand before terminal differentiation.

Definitive HSC and their progeny are dependent on the transcription factor *c-Myb* whereas primitive YS-derived erythrocytes, megakaryocytes (Tober *et al.*, 2008) and tissue-resident macrophages (Schulz *et al.*, 2012) are *c-Myb* independent. Ter119^+^ cells were drastically decreased in the FL of *c-Myb* mutants and P1 cells were undetectable. By comparison, the CD45^+^ cells present in the FL of *c-Myb* mutants representing tissue resident macrophages were not affected (Schulz *et al.*, 2012). Ter119^+^ cells in the *c-Myb*^-/-^ FL expressed embryonic globins (εy and βh1), consistent with their primitive origin and do not express *Klf1*, a transcriptional activator of the β-globin promoter essential for the transition from expression of embryonic to adult hemoglobin (Perkins *et al.*, 1995, 2016). Erythroid transcripts were not detected in *c-Myb*^-/-^ TER119^-^CD24^+^ or CD24^-^ cells indicating that no definitive erythropoiesis developed in these mutants. Previous studies reported that *c-Myb* mutations affected both embryonic erythropoiesis and HSC progeny, which was taken as evidence for the HSC origin of embryonic erythrocytes. However, c-Myb induces proliferation of erythroid progenitors and therefore HSC-independent erythroid cells can also be affected by *c-Myb* mutations (Vegiopoulos *et al.*, 2006).

It was previously shown that YS-derived EMP transplanted in adult recipient mice persist for around 20 days reflecting the half-life of erythrocytes *in vivo* (McGrath *et al.*, 2015a). These experiments however do not evaluate the relative contribution of EMPs and HSCs to erythropoiesis in an unperturbed environment.

Single injections of OH-TAM at E8.5 in Csf1r^MeriCreMer^Rosa26^YFP^ mark exclusively YS-derived cells and their progeny, among which is the microglia (Gomez Perdiguero *et al.*, 2015). In these mice P1 and P2 cells are marked at levels similar to the microglia, three days after OH-TAM, indicating they are the progeny of YS EMPs. Consistent with the lineage relationship previously established, the frequency of labelled P1/P2 cells decreased with time after injection whereas the frequency of labelled erythrocytes increased. EMPs emerge in the YS between E8.5-10.5 and an injection of OH-TAM at E8.5 will lead to the highest circulating levels of the drug 6 hours later, rapidly decreasing thereafter (Zovein *et al.*, 2008) leading to the labelling of only a fraction of EMP. EMPs differentiating into any myeloid progeny at the time of injection will maintain expression of Csf1r and will be labelled with YFP. By contrast, differentiation into erythroid progenitors that expand and progress to erythroblasts lose Csf1r expression throughout, thus explaining the decreasing frequency of labelled immature erythroid progenitors with time. We have demonstrated by intraembryonic injections that P2 cells rapidly differentiate into Ter119^+^ circulating RBC.

A kinetics that mirror images the one described above is found in Flt3^Cre^Rosa26^YFP^ mice where Flt3 expressing cells and their progeny are permanently labelled with YFP. By E16.5, where equivalent frequencies of LSK, CMP, GMP and CD11b^+^ monocytes are YFP^+^, only a small frequency of erythroid progenitors including MEP and virtually undetectable frequencies of P3 or erythroblasts are labelled. By E18.5 MEP were labelled at similar levels to those of CMP and monocytes although the erythroblast compartment still shows a modest contribution of Flt3 expressing progenitors. Because lymphoid progenitors persistently express Flt3 after commitment they are the highest labelled population in this model and were excluded from the analysis. It has been recently described that megakaryocyte/erythrocyte progenitors can bypass the stage of Flt3 expressing MPP and would therefore be undetected in this model, although no FL counterpart has been reported (Carrelha *et al.*, 2018). However, less than 5% of CD150^+^CD48^-^HSC appear to adopt this behavior, a low frequency that will not impact our conclusions. In addition, in adult Flt3-Cre YFP mice the frequencies of YFP labelled erythroid progenitors is similar to that of all other HSC-derived lineages indicating this is a suitable model to analyze the erythroid lineage (Boyer *et al.*, 2011; Buza-Vidas *et al.*, 2011; Gomez Perdiguero *et al.*, 2015).

HSC in FL expand but also differentiate giving rise to multilineage progeny that comprise CMP, GMP, and lymphoid progenitors. However, our data shows that despite a rapid differentiation of FL HSC, they do not contribute significantly to the erythroid compartment before birth and therefore *in vivo* embryonic HSC differentiation is skewed (Supplementary Fig. 5). The FL stromal microenvironment can sustain erythropoiesis and FL HSC can differentiate into erythrocytes *in vitro*. We show that the low levels of Epo available in FL before the kidney is competent to produce adult levels of this hormone modulate the differentiation pattern of HSC that do not produce MEP and do not contribute to erythropoiesis (Zanjani *et al.*, 1981). Large numbers of expanding YS-derived erythrocyte progenitors efficiently outcompete HSC progeny in an environment where resources for erythroid differentiation are limiting.

These results reinforce the notion that in contrast to what has been accepted, YS hematopoiesis is not only devoted to providing oxygen to the embryo before HSCs differentiate in FL (Supplementary Fig. 5) but rather sustain embryonic survival until birth. A recent report analyzing human fetal liver hematopoiesis indicates that all cells in the erythrocyte lineage, similar to the observation in the mouse reported here, do not express CD45 at stages ranging from 7-17 weeks post conception (Popescu *et al.*, 2019). These observations suggest that fetal erythropoiesis originates in the YS, also in humans and will impact our understanding of embryonic hematopoiesis in general and in the pathogenesis of infant erythrocyte abnormalities.

## Materials and Methods

### Mice

C57BL/6J mice were purchased from Envigo, Ubiquitin–GFP (Schaefer *et al.*, 2001) mice used for transplantation experiments were a kind gift from P. Bousso (Pasteur Institute) Myb^-/-^, Csf1r^MeriCreMer^, Flt3^Cre^ and Rosa26^YFP^ mice have been previously described (Gomez Perdiguero *et al.*, 2015). 6-8-week-old mice or timed pregnant females were used. Timed-pregnancies were generated after overnight mating, the following morning females with vaginal plug were considered to be at E0.5. Recombination in Csf1r^MeriCreMer^Rosa26^YFP^ was induced by single injection at E8.5 of 75 µg per g (body weight) of 4-hydroxytamoxifen OH-TAM (Sigma), supplemented with 37.5 µg per g (body weight) progesterone (Sigma) as described (Gomez Perdiguero *et al.*, 2015). All animal manipulations were performed according to the ethic charter approved by French Agriculture ministry and to the European Parliament Directive 2010/63/EU.

### Cell suspension

E11.5-E18.5 fetal livers (FL) were dissected under a binocular magnifying lens. FLs were recovered in Hanks’ balanced-salt solution (HBSS) supplemented with 1% fetal calf serum (FCS) (Gibco) and passed through a 26-gauge needle of a 1-ml syringe to obtain single-cell suspensions. Before staining, cell suspensions were filtered with a 100 μm cell strainer (BD).

### Flow cytometry and cell sorting

For sorting, FL were depleted (Ter119^+^CD45^+^) using MACS Columns (Miltenyi Biotec). Cell suspensions were stained for 20-30 min at 4°C in the dark with antibodies listed in Supplementary Table 2. Biotinylated antibodies were detected by incubation for 15 min at 4°C in the dark with streptavidin. Antibodies to lineage markers included anti-Ter119, anti-Gr1, anti-CD19, anti-CD3, anti-CD4, and anti-CD8, anti-NK1.1, anti-Il7r, anti-TCRαβ, anti-TCRγδ and anti-CD11c (all identified in Supplementary Table 2). Stained cells were analyzed on a custom BD LSR Fortessa or BD FACSymphony and were sorted with a BD FACSAria III (BD Biosciences) according to the guidelines for the use of flow cytometry and cell sorting (Cossarizza *et al.*, 2019). Data were analyzed with FlowJo software (v.10.5.3, BD Biosciences) or R packages as described in “Bioinformatic Analysis”.

### RNA-sequencing and analysis

Total RNA from sorted E14.5 FL cells was extracted using the RNeasy Micro kit (Qiagen) following manufacturer instructions and rRNA sequences were eliminated by enzymatic treatment (Zap R, Clontech). cDNA libraries were generated using the SMARTer Stranded Total RNA-Seq Kit – Pico Input Mammalian (Clontech). The single read RNA-seq reads were aligned to the mouse reference genome GRCm38 using STAR. Number of reads aligned to genes were counted using FeatureCounts (Liao *et al.*, 2014). The R package DESeq2 (Love *et al.*, 2014) was used to normalize reads and identify differentially expressed genes with statistically significance using the negative binomial test (p<0.05, Benjamini-Hochberg correction).

Enrichr was used to perform gene-set enrichment analysis of the highly differentially expressed genes in P2 vs CD324^+^ cells (>2-fold differential expression) (Chen *et al.*, 2013). Top 10 terms from the Gene Ontology Biological Process 2018 and ARCHS4 Tissues were retrieved. Expression datasets are available in NCBI Gene Expression Omnibus under Accession Number GSE138960.

### Gene expression by RT-PCR

Cells were sorted directly into lysis buffer and mRNA was extracted (RNeasy Micro Kit (Qiagen), reverse-transcribed (PrimeScript RT Reagent Kit (Takara Bio), and quantitative PCR with Power SYBR Green PCR Master Mix (Applied Biosystems)(see Supplementary Table 3 for primers). qPCR reactions were performed on a Quantstudio3 thermocycler (Applied Biosystems), gene expression was normalized to that of β-actin and relative expression was calculated using the 2^−ΔCt^ method.

### Imaging Flow Cytometry Analysis

E13.5 FL cells were stained with the surface markers CD45 BV605 (1:50 dilution), Ter119 Biotin (1:100 dilution) followed by incubation with Streptavidin PE-Cy7 (1:100 dilution), CD71 PE (1:100 dilution), CD24 BV510 (1:50 dilution) and Kit Pacific Blue (1:20 dilution) and the RNA Dye Thiazole Orange (TO). Prior to acquisition, nuclei were stained with 20 μM DRAQ5 (Biostatus) and filtered with 100 μm mesh. Data acquisition was performed using an ImageStream^x^ Mark II Imaging Flow Cytometer (Amnis, Luminex Corporation) using 405 nm, 488 nm, 561 nm and 642 nm excitation lasers and the 40× magnification collection optic. Laser powers were set in order to maximize signal resolution but minimize any saturation of the CCD camera with bright-field images collected in channels 1 and 9. A minimum of 100,000 cell events were collected per sample. In order to calculate spectral compensation, single-stained cells were acquired with the bright-field illumination turned off. Spectral compensation and data analysis were performed using the IDEAS analysis software (v.6.2.64, Luminex Corp).

### EdU incorporation and cell cycle analysis

EdU detection was done using the Click-iT EdU pacific blue flow cytometry assay kit (Invitrogen). Cell cycle was analyzed after fixation with Fixation/Permeabilization kit (eBioscience™) and staining with Ki67. DAPI (4,6-diamidino-2-phenylindole) was added 7 min before analysis.

### *In vitro* liquid and semi-solid cultures

For limiting dilution analysis sorted cells were plated in 1:3 diluting densities starting at 27 cells/well until 1 cell/well in complete medium OPTI-MEM with 10% FCS, penicillin (50 units/ml), streptomycin (50 μg/ml) and β-mercaptoethanol (50 μM) supplemented with a saturating amount of the following cytokines: macrophage colony-stimulating factor (M-CSF), granulocyte-macrophage colony-stimulating factor (GM-CSF), c-Kit ligand (Kitl), Erythropoietin (Epo) (R&D Systems) and Thrombopoietin (Tpo) (R&D Systems) for myeloid and erythroid differentiation. Except stated otherwise, cytokines were obtained from the supernatant of myeloma cell lines (provided by F. Melchers) transfected with cDNA encoding those cytokines. After 5-7 days wells were assessed for the presence of hematopoietic colonies. Cell frequencies, determined with ELDA software (‘extreme limiting-dilution analysis’) from the Walter and Eliza Hall Institute Bioinformatics Division, are presented as the number of positive wells and the number of total tested wells (Hu and Smyth, 2009). For colony forming assays sorted cells were plated at 100 cells/35 mm culture dishes in duplicates in Methocult M3434 as described by the manufacturer. CFU-E were assessed at 3 days and remaining colonies at 7 days.

### *In vivo* analysis of lineage potential

E13.5 GFP^+^ Kit^+^CD24^+^ or CD45^+^LK FL cells from UBC-GFP embryos were purified and injected intraperitoneally into recipient E13.5 WT embryos (20000 cells/embryo) of anesthetized females and FL and fetal blood were analyzed at E16.5.

### Multiplex single-cell qPCR

Single cells were sorted directly into 96-well plates loaded with RT-STA Reaction mix (CellsDirect^TM^ One-Step qRTPCR Kit, Invitrogen, according to the manufacturer’s procedures) and 0.2x specific TaqMan® Assay mix (see Supplementary Table 4 for the TaqMan® assays list) and were kept at −80 °C at least overnight. For each subset analyzed, a control-well with 20 cells was also sorted. Pre-amplified cDNA (20 cycles) was obtained according to manufacturer’s note and was diluted 1:5 in TE buffer for qPCR. Multiplex qPCR was performed using the microfluidics Biomark HD system for 40 cycles (Fluidigm) as previously described (Chea *et al.*, 2016). The same TaqMan probes were used for both RT/pre-amp and qPCR. Only single cells for which at least 2 housekeeping genes could be detected before 20 cycles were included in the analysis.

### Bioinformatic Analysis

Flow cytometry data analysis was performed in FCS files of live CD45-Ter119-cell fraction using R packages “Rtsne”, “Rphenograph” and “pheatmap” using Rv3.5.0. Gene expression raw data (BioMarkTM, Fluidigm) of single cells was normalized with Gapdh and β-actin. Heatmaps and hierarchical clustering were generated using R packages “pheatmap” and “Rphenograph” (Levine *et al.*, 2015).

### Quantification and Statistical Analysis

All results are shown as mean ± standard deviation (SD). Statistical significance was determined using one-way ANOVA followed by Tukey multiple comparison test where a P value of <0.05 was considered significant and a P value >0.05 was considered not significant.

## Acknowledgements

We thank J. Palis, A. Bandeira, P. Vieira and P. Pereira and the members of P.P.O. and A.C. laboratories for critical discussions, M. Petit for help in bioinformatic analysis. We thank P. Bousso for the ubiquitin-GFP reporter mice; S. Novault, S. Megharba, S. Schmutz, V. Libri and T. Stephen from the Center for Translational Science (CRT) – Cytometry and Biomarkers Unit of Technology and Service (CB UTechS) at Institut Pasteur for technical support; the staff of the animal facility of the Institut Pasteur for mouse care. This work was financed by the Institut Pasteur, INSERM, ANR (grant Twothyme) to A.C., REVIVE Future Investment Program and Pasteur-Weizmann Foundation through grants to A.C., FCT through the grant “PD/BD/114128/2015” to F.S.S and the grant “FCT-POCI-01-0145-FEDER-032656” to P.P.O..

## Author Contributions

F.S.S.: Conceived and performed experiments, performed formal analysis, wrote the manuscript

O. B-D.: Performed experiments

R. E.: Performed experiments and provided feedback

L. I.: Performed experiments

L. F.: Performed experiments

O. S.: Performed RNA-Seq

P. P.Ó.: provided feedback and reviewed the manuscript

E. G-P.: provided animals, expertise and feedback, reviewed the manuscript

A. C.: Conceived and performed experiments, performed formal analysis, wrote and reviewed the manuscript, secured funding

All authors discussed and interpreted the results.

## Data availability

The accession number for the RNA-seq data reported in this paper is GSE138960. For original data, please contact

## Supplementary Figure Legends

**Supplementary Figure 1.**
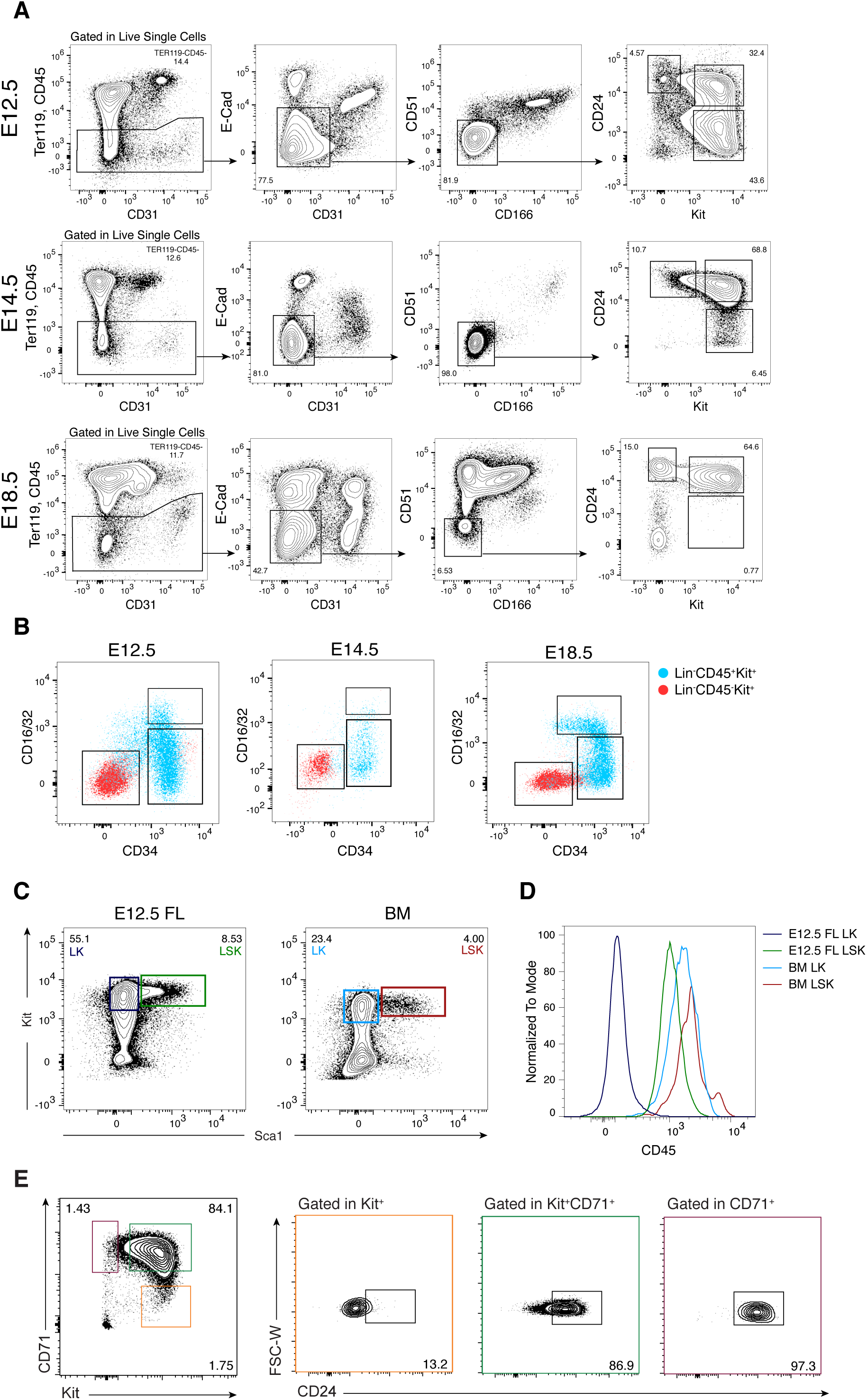
Phenotype of E12.5, E14.5 and E18.5 fetal liver and adult bone marrow populations (related to Figure 1 and 3) **(A)** Flow cytometry analysis of E12.5, E14.5 and E18.5 fetal liver cells using Ter119, CD45, E-Cadherin, CD31, CD51 and CD166. Viable Ter119^-^CD45^-^E-Cad^-^CD31^-^ CD51^-^CD166^-^ cells can be subdivided into 3 populations according to expression of CD24 and Kit. **(B)** Flow cytometry profile of E12.5, E14.5 and E18.5 fetal liver Lin^-^ CD45^+^Kit^+^ (CD45^+^LK) (blue) and CD45^-^Kit^+^ (red) cells according to CD16/32 and CD34 expression. **(C)** Flow cytometry analysis of CD24 expression in P3 (CD71^+^Kit^-^), P2 (CD71^+^Kit^+^) and P1 (CD71^-^Kit^+^) cells in E13.5 FL cells.

**Supplementary Figure 2.**
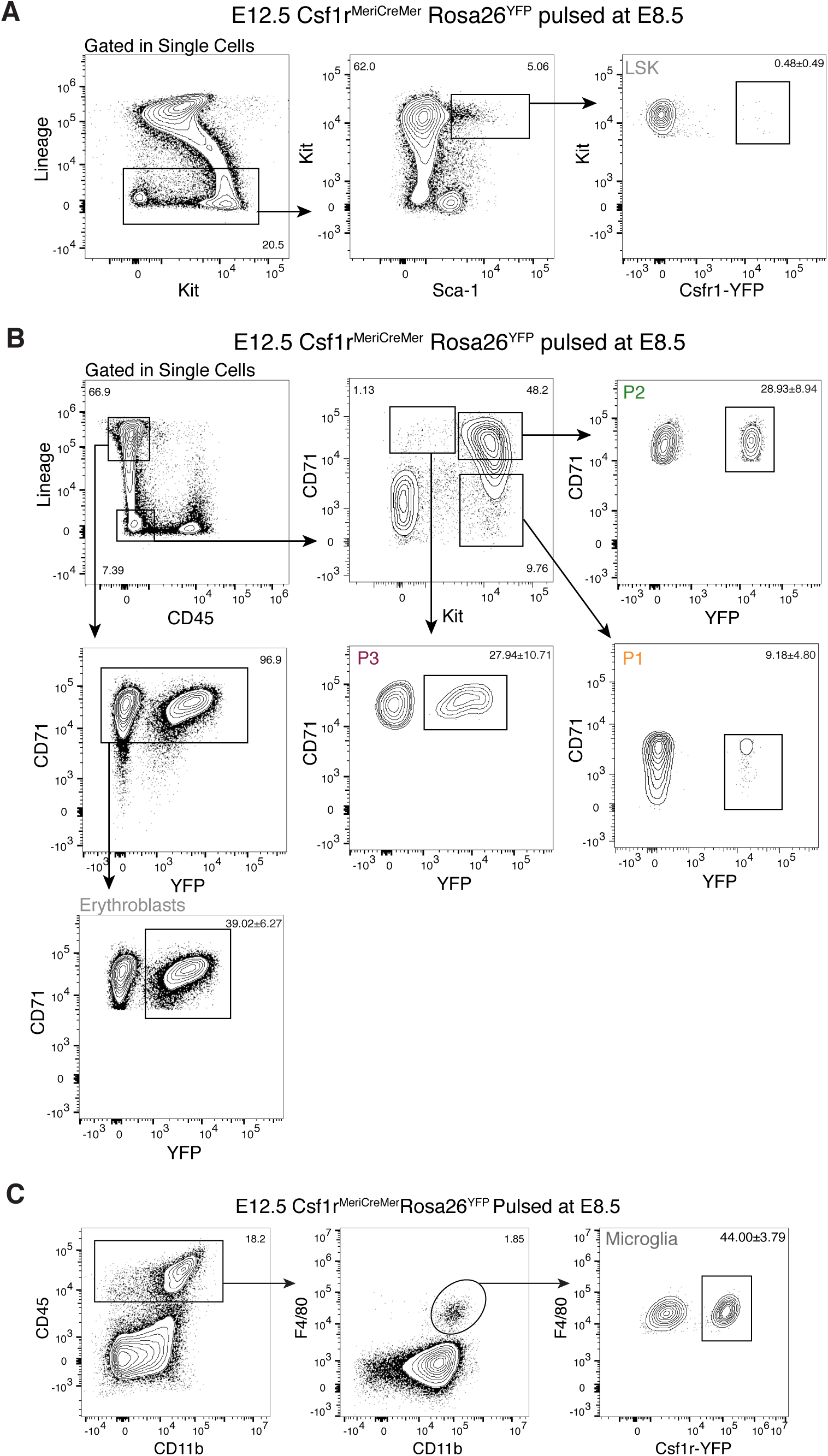
Gating strategies used to determine YFP labelled hematopoietic populations in the Csf1r^MeriCreMer^Rosa26^YFP^ lineage tracing model (related to Figure 6) **(A)** Gating strategy for the analysis of YFP expression in Lin^-^Kit^+^Sca1^+^ (LSK) cells. **(B)** Gating strategy for the analysis of YFP expression in P1 (Lin^-^CD45^-^Kit^+^CD71^-^) P2 (Lin^-^CD45^-^Kit^+^CD71^+^) cells and P3 (Lin^-^CD45^-^Kit^-^CD71^+^) and erythroblast (Lin^+^CD71^+^) cells. **(C)** Gating strategy for the analysis of YFP expression in microglia (CD45^+^ F4/80^+^ CD11b^+^) cells. Data representative of E12.5 YFP^+^ embryos.

**Supplementary Figure 3.**
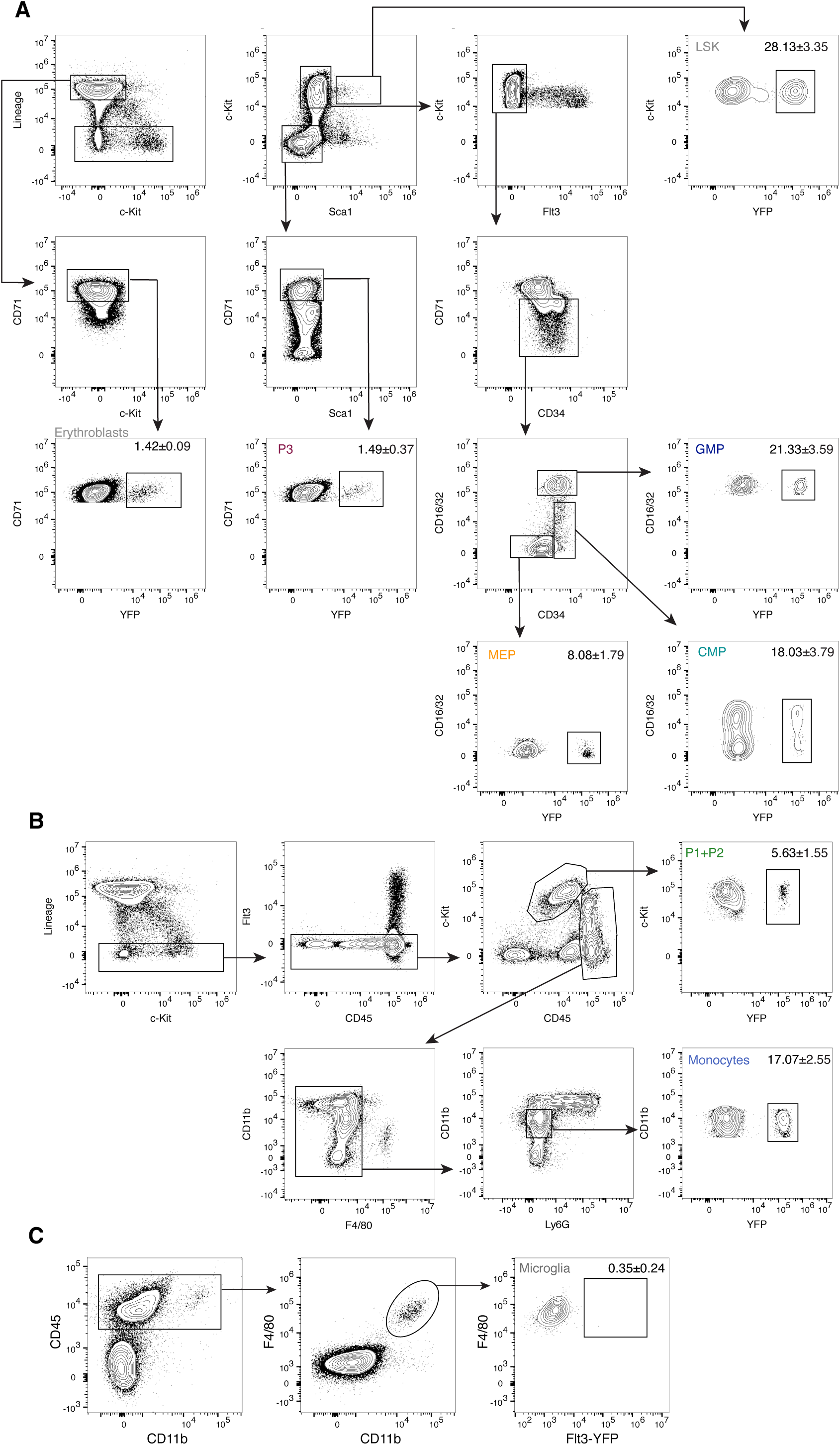
Gating strategies used to determine YFP labelled hematopoietic populations in the Flt3^Cre^Rosa26^YFP^ fate-mapping model (related to Figure 6) **(A)** Gating strategy for the analysis of YFP expression in Lin^-^Kit^+^Sca1^+^ (LSK), Common Myeloid Progenitors (CMP) (Lin^-^Kit^+^Sca1^-^Flt3^-^CD71^-^CD16/32^-^CD34^+^), Granulocyte-Monocyte Progenitors (GMP) (Lin^-^Kit^+^Sca1^-^Flt3^-^ CD71^-^ CD16/32^+^CD34^+^), Megakaryocyte Erythrocye Progenitors (MEP) (Lin^-^Kit^+^Sca1^-^Flt3^-^ CD71^-^CD16/32^-^CD34^-^), P3 (Lin^-^Kit^-^CD71^+^) and erythroblast (Lin^+^CD71^+^) cells. **(B)** Gating strategy for the analysis of YFP expression in P1+P2 (Lin^-^CD45^-^Kit^+^) and monocytes (Lin^-^CD45^+^CD11b^+^F4/80^-^) cells. **(C)** Gating strategy for the analysis of YFP expression in microglia (CD45^+^ F4/80^+^ CD11b^+^) cells. Data representative of E16.5 YFP^+^ embryos.

**Supplementary Figure 4.**
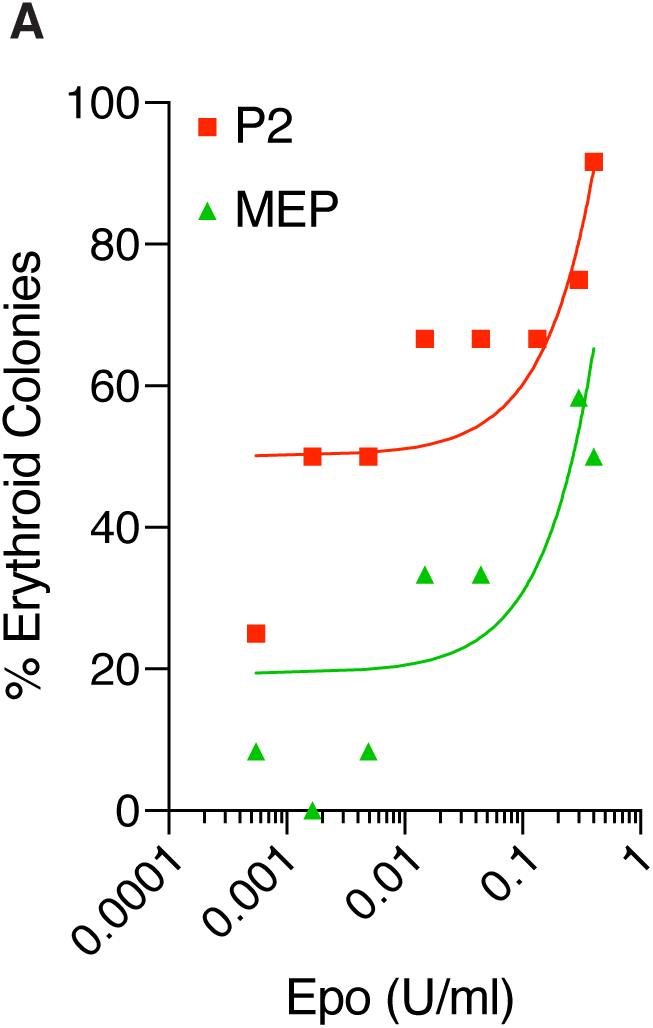
Erythroid colony formation in response to erythropoietin **(A)** Frequency of erythroid colonies in P2 and MEP (CD45^+^) cells from E13.5 FL using serial dilutions of erythropoietin. Representative plot of 3 independent experiments. Curves represent the linear regression of the data.

**Supplementary Figure 5.**
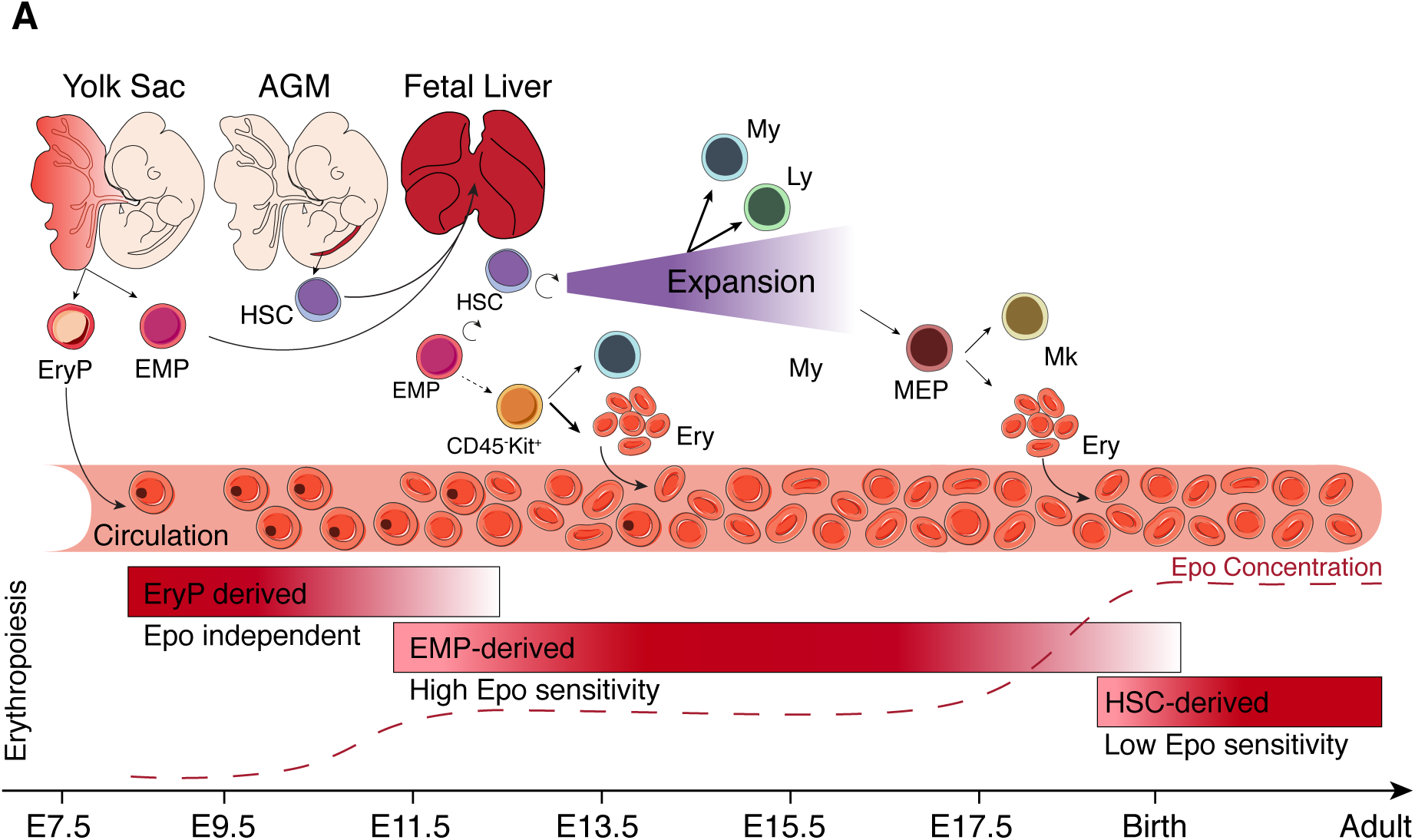
Model of embryonic erythropoiesis **(A)** Erythroid progenitors (P-Ery) are first generated in the yolk sac (E7.5) and give rise to primitive nucleated erythrocytes, still found in low frequencies at birth. A second wave of progenitors emerges in the yolk sac (E8.5) as erythromyeloid progenitor (EMP). EMPs migrate to the FL where they differentiate into highly proliferative erythroid (CD45^-^ Kit^+^) progenitors that sustain erythropoiesis during embryonic life. Hematopoietic stem cells (HSC) generated in the AGM (E9.5-E11.5) migrate to the FL where they expand and differentiate into myeloid and lymphoid lineages. Contribution of HSC to the erythroid lineage is only detected after birth. YS-derived progenitors respond to lower levels of erythropoietin than their HSC counterparts and have a selective advantage in FL where erythropoietin levels are lower than in adult BM. Abbreviations: YS: yolk sac; AGM: aorta-gonads-mesonephros; FL: Fetal Liver; P-Ery: Primitive Erythroid Progenitors; EMP: Erythromyeloid Progenitors; HSC: Hematopoietic Stem Cells; Ly: Lymphoid Lineages; My: Myeloid Lineages; Ery: Erythroid Cells; MEP: Megakaryocyte/Erythrocyte Progenitors.

## Supplementary Tables

**Supplementary Table 1.**
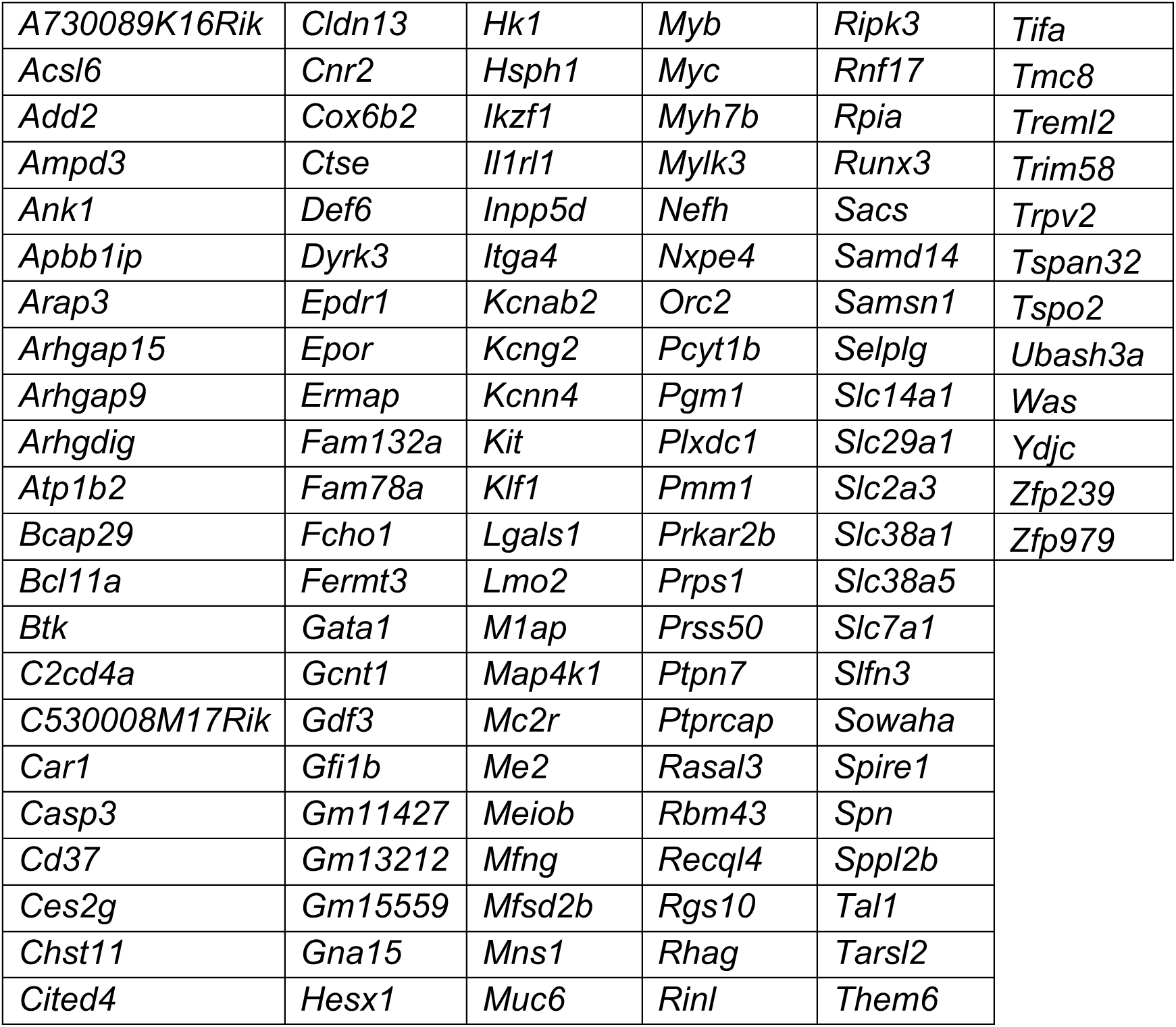
List of the 122 genes submitted to Enrichr, related to Figure 2

**Supplementary Table 2.**
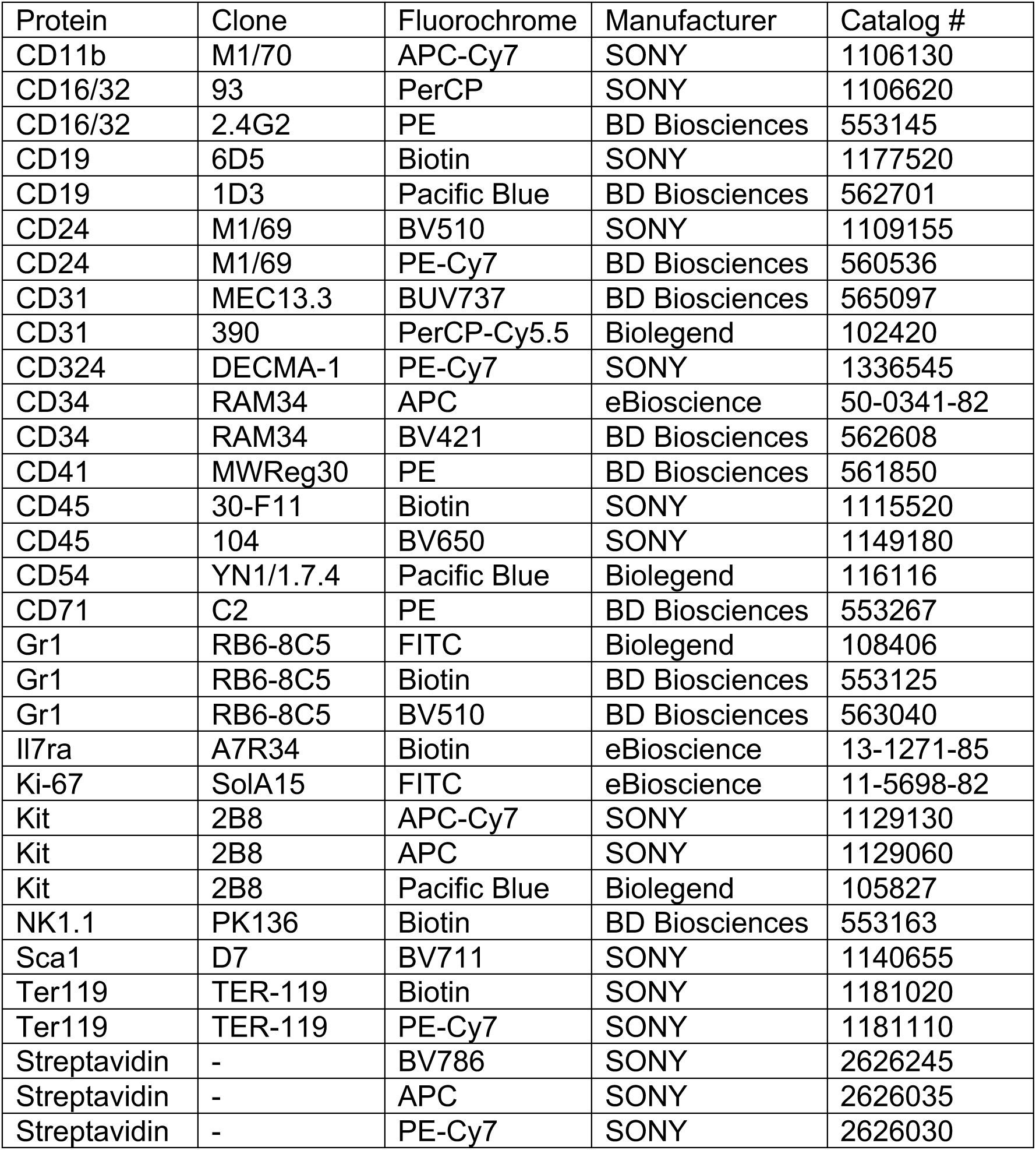
List of antibodies for flow cytometry

**Supplementary Table 3.**
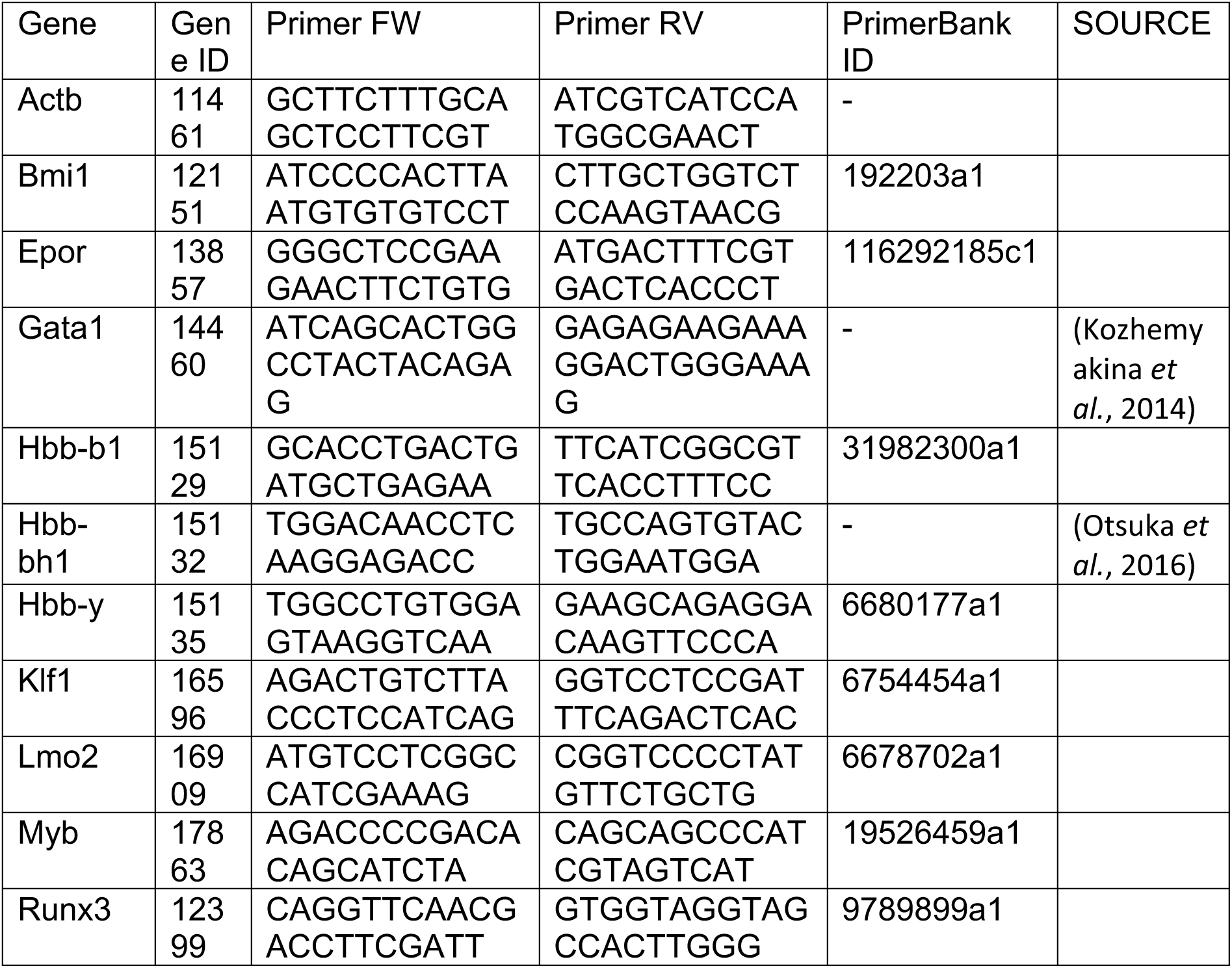
List of Primers for qRT-PCR

**Supplementary Table 4.**
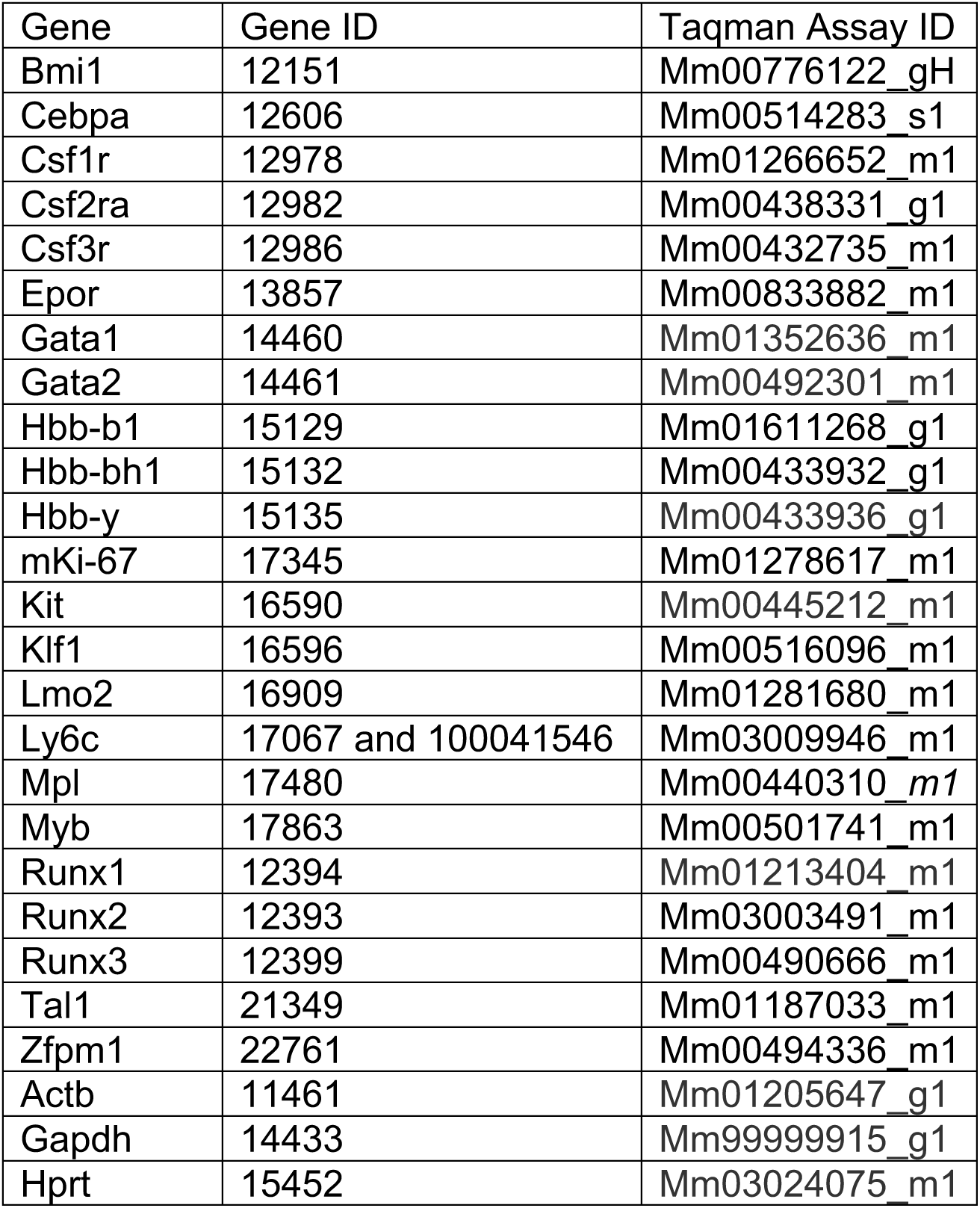
Taqman probes for Single-Cell Multiplex Gene Expression

